# AlphaFold Models Illuminate Half of Dark Human Proteins

**DOI:** 10.1101/2021.11.04.467322

**Authors:** Jessica L. Binder, Joel Berendzen, Amy O. Stevens, Yi He, Jian Wang, Nikolay V. Dokholyan, Tudor I. Oprea

## Abstract

We investigate the use of confidence scores to predict the accuracy of a given AlphaFold model for drug discovery. Predicted accuracy is improved by eliminating confidence scores below 80, due to effects of disorder. 95% of models corresponding to a set of recent crystal structures are accurate at the fold level. Conformational discordance in the training set has a more significant effect on accuracy than sequence divergence. We propose criteria for models and residues that are possibly useful for virtual screening, by which AlphaFold provides models for half of understudied (dark) human proteins and two-thirds of residues in those models.

## INTRODUCTION

About half of Americans answering a 2020 survey would not get in an AI-driven taxi, and about three-quarters of them believed AI (artificial intelligence) cars were “not ready for primetime” [1]. Whether driving a vehicle or discovering new medicines, trust in AI depends on accumulated community experience and the consequences of errors in specific cases. There are over 20,000 protein-coding genes in the human genome [2–4]. Of these, 7074 have experimentally determined structures deposited in the Protein Data Bank (PDB) as of July 2021 [5]. Only 670 human proteins are therapeutically targeted by medicines, comprising the “drugged genome” [6]. Significant areas of biology remain potentially amenable to drug discovery [7]. Initiatives like “Illuminating the Druggable Genome” [8], OpenTargets [9], and Target 2035 [10] are exploring novel therapeutic opportunities in the “druggable” genome.

DeepMind described [11,12] AF2 (AlphaFold version 2.0), an AI method that predicts overall 3D structures of proteins. More than 350,000 AF2 structural models (including models of nearly every human protein) are now publicly accessible [12]. DeepMind garnered worldwide attention with their decisive win of the Critical Assessment of Techniques for Protein Structure Prediction, CASP14 [13]. Currently, scientists are assessing the impact of AF2 on research, including how much AF2 models expand the druggable genome.

Winning CASP14 presents a set of challenges specific to protein folding. However, protein 3D models do not often play a crucial role in drug discovery. The notion of *trust* in AF2 models is illustrated with a histogram of atomic Root-Mean-Square Deviations (*aRMSD*) on CIZ atoms for crystal structures deposited in the PDB since AF2 was trained (*Fig. 2a* in [11]). It shows that AF2 produces high-quality folds in two-thirds of cases. However, the overall accuracy of a given AF2 model was not discussed. Local confidence scores (predicted Local Distance Difference Test, *pLDDT*) show a 95% per-residue correlation with experimentally-derived *LDDT* values [14] over the same proteins. AF2 model confidence evaluation is needed in the drug discovery context, given the non-local nature of *aRMSD*, the inherent selection bias of recent PDB structures, and the lack of any overall confidence-in-accuracy measure that can be calculated for individual models. Here, we discuss the issue of trust in AF2 models by addressing disorder, divergence, discordance, and druggability.

### Disorder dominates confidence scores below 80

More than 30% of eukaryotic proteins contain one or more intrinsically disordered regions, IDRs [15– 21]. Disorder is reflected in confidence scores as regions with low *pLDDT* [12]. **Figure 1** displays the distributions of the *pLDDT* scores reported by AF2 for resolved/ordered and unresolved/disordered regions of crystal structures deposited in the PDB since AF2 was trained (the “post-AF2 test set”, see **Supplemental Information**). On this set of structures, ordered regions most frequently show *pLDDT* scores greater than 80, while IDRs have a broad distribution of *pLDDT* scores, with about 40% of unresolved regions falling below a *pLDDT* score of 50. From this analysis, we conclude that pLDDT confidence scores below 80 are more indicative of disorder than of confidence in the accuracy of ordered structures.

**Figure 1.**
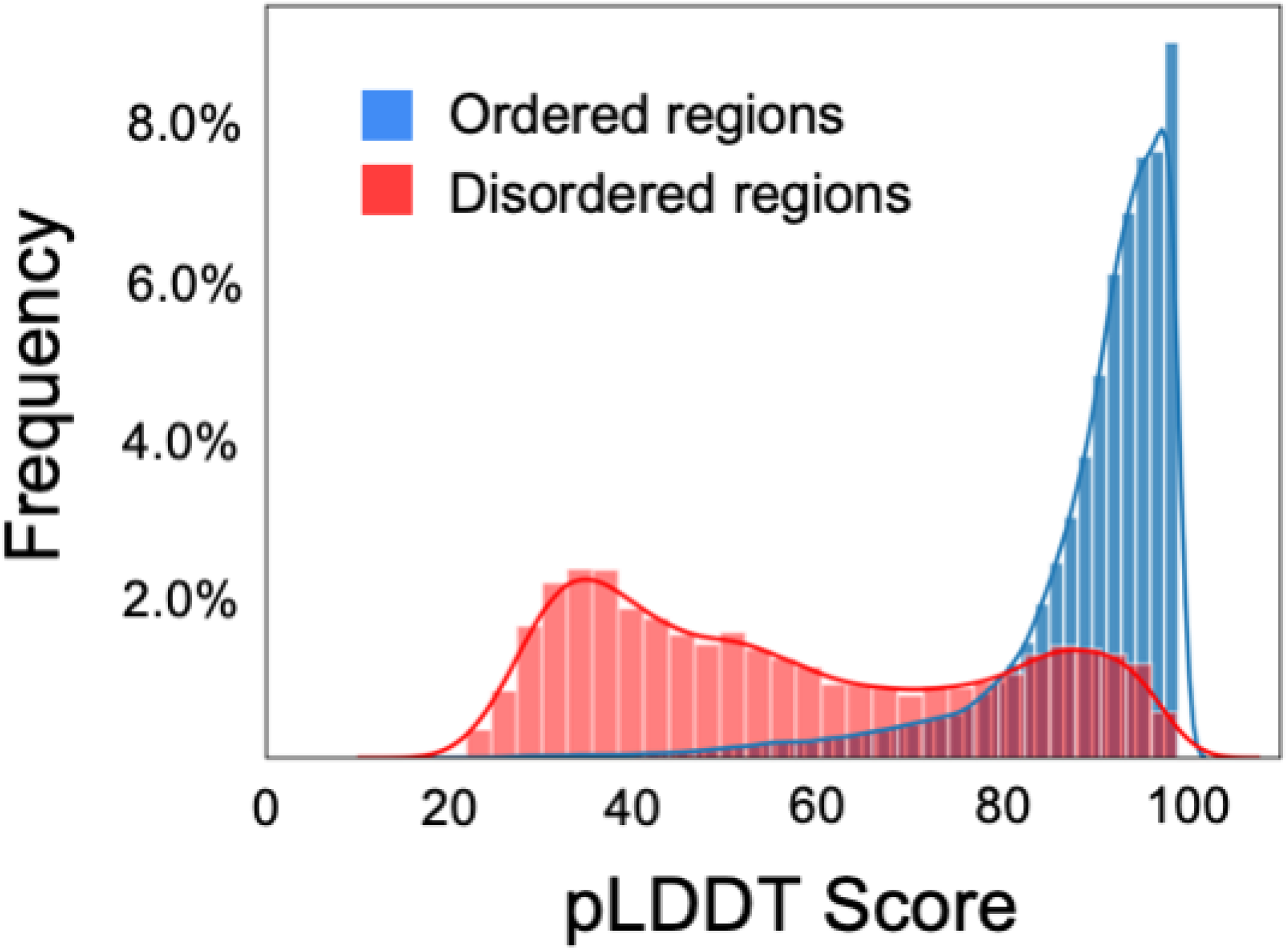
Distribution of AlphaFold confidence scores across ordered (blue) and disordered (red) regions. Ordered and disordered regions correspond to resolved and unresolved parts, respectively, for the post-AF2 test set. Terminal regions were not included. Ordered regions most frequently show pLDDT scores >80%. Disordered regions show a broad distribution of pLDDT scores with comparable frequencies from pLDDT scores between 20% and 90%.

### Divergence has a minor effect on model accuracy

A problem with the 6-bin histogram used to estimate the distribution of model accuracies [11] is that *aRMSD* is a non-local measure. If a model is incorrect at the fold level, the expectation value of *aRMSD* scales with the radius of gyration. Thus, a model with 30 Å *aRMSD* against the experimental structure could be consistent with an entirely misfolded domain of around 1000 residues in length [22] or simply with rotation of a smaller domain about a single residue. To characterize different effects on model accuracy, we down-selected the post-AF2 test set to 1,779 models that can be aligned with a corresponding experimental structure (see **Supplemental Information**) and used them to evaluate all-atom and backbone measures. *pLDDT* correlates poorly with log(*aRMSD*) on this set: Spearman rank correlation coefficient is 0.43 on the median (**Supplemental Figure S1A)**. Truncating the range of *pLDDT* over which the median is calculated with a floor of 80 slightly improves the coefficient to −0.48 (**Supplemental Figure S1B**). We refer to the per-model median value of *pLDDT* scores greater than 80 as *pLDDT*_*80*_.

Next, we split this down-selected test set into two pairs of subsets. The first pair explored high (*pLDDT*_*80*_ > 90) or low (*pLDDT*_*80*_ < 88) confidence scores. The second pair explored high (in clusters at 100% identity for over 80% of the length) or low (out of clusters at 5% identity for over 80% of the length) sequence identity to structures previously in the PDB. Cutoff values in these pairings were chosen to give maximal differences while maintaining roughly comparable fractions of the test set in the two arms of each pairing. We calculated distributions on log(*aRMSD*) and on the all-atom *LDDT* [14] for each of the subsets (**Figure 2**).

**Figure 2.**
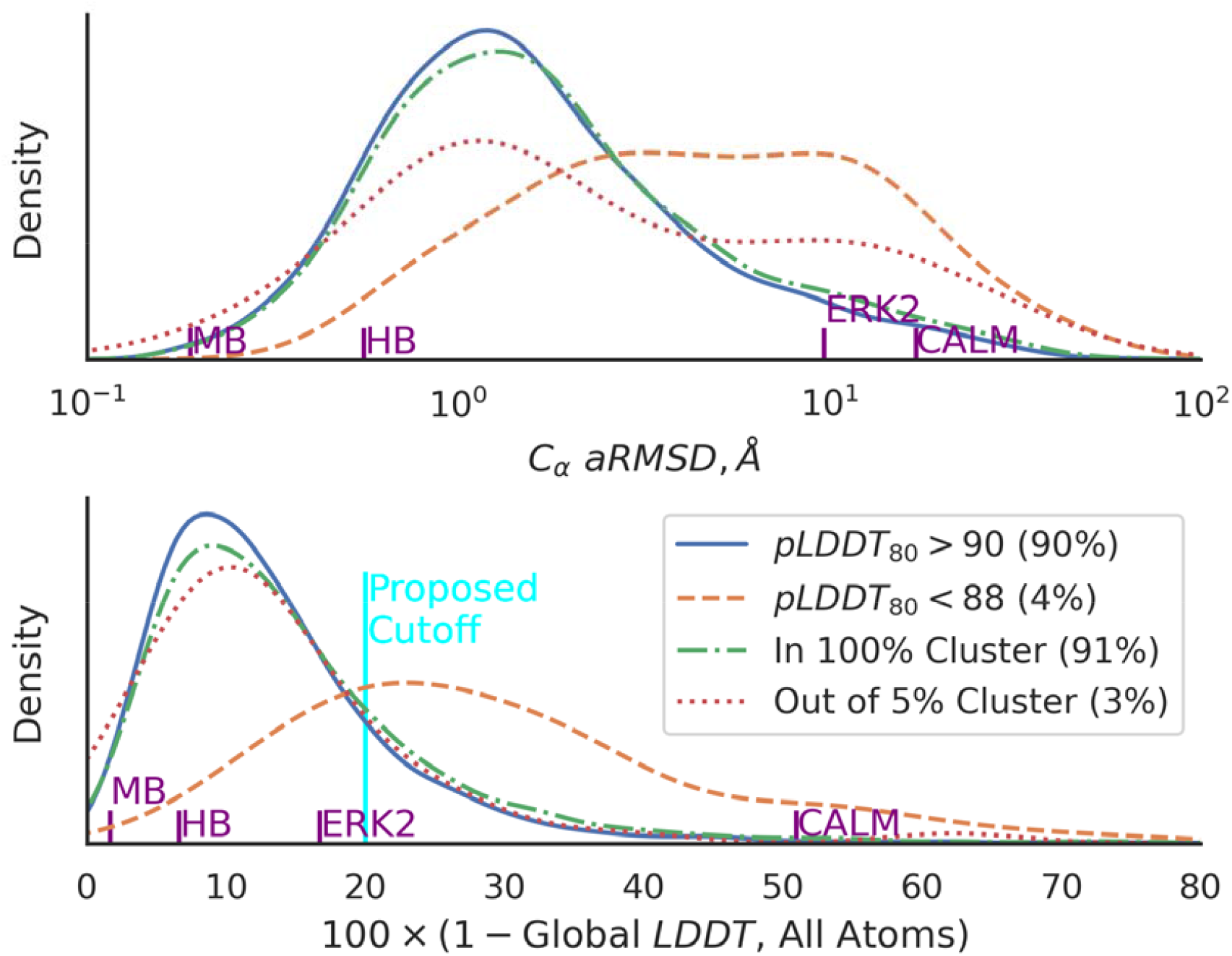
Kernel-density estimates of the distribution of differences between AF2 models and crystal structures using atomic Root-Mean-Square Displacements on Cα atoms (top) and Global Local Distance Difference Test metrics (bottom). These distributions were calculated from crystallographic structures that were deposited in the Protein Data Bank after the AF2 training set cut-off date of April 30, 2018. Only residues that unambiguously intersect between AF2 models deposited in EMBL [47] and crystal structures were considered, with a minimum per-chain length cutoff of 20 residues, resulting in 1810 structural models to be compared. Distributions for mean confidence levels (pLDDT_80_) over the raw models at or above 90 (solid blue line) and below 88 (dotted orange line) are plotted. We also clustered the PDB using mmseqs [48] to select for sequences nearly identical to an existing structure (in clusters with 100% minimum sequence identity over 80% of the longest sequence and cluster mode 2, dot-dash green line) or decisively non-matching regions (out of 5% minimum sequence identity, dotted red line). The high-confidence (pLDDT_80_>90) distribution on log(aRMSD) peaks at 1.7 □ aRMSD, with a long tail extending beyond 10 Å at the 10% level. The low-confidence distribution on log(aRMSD) has a broad flat shape suggesting peaks at 3 and 20 Å. The high-identity distribution looks similar to the high-confidence distribution, while the low-identity distribution has peaks near 2 and 20 Å, respectively. Plotted against all-atom LDDT, the high-confidence, high-identity, and low-identity distributions look similar to each other; only the low-confidence distribution is distinct, with a single peak at LDDT ∼75. Notations on the x-axis indicate differences between structures of ligand-free vs. ligand bound myoglobin (MB, PDB entries 1A6N and 1A6G); R-vs. T-state hemoglobin (HB, PDB entries 6BWP and 6BWU); unphosphorylated vs. doubly-phosphorylated conformations of an extracellular signal-regulated kinase (ERK2, PDB entries 1ERK and 2ERK0); and calcium-free vs. calcium-bound calmodulin (CALM, PDB entries 1CLL and 1QX5). The cyan line shows the proposed LDDT cutoff for a structure that is likely to be useful for virtual screening.

The *aRMSD* metric is not well-suited for characterizing structural models because its non-local nature tends to exaggerate the effects in small number backbone angle changes [23]. The lack of a high-difference in the low-identity *LDDT* distribution, together with the suppression of the high-difference tail in the low-confidence distribution, suggest that most differences between model and structure are in a few local coordinates, rather than many. Using *LDDT* as the accuracy measure improved Spearman’s correlation on *pLDDT*_80_ to 0.60 (**Supplemental Figure S2)**.

Less than 1% of the high-confidence distribution appears in the range consistent with fold-level inaccuracies at *LDDT<*50. Distributions for the low-similarity, high-confidence, and high-identity subsets are the same, with the caveat afforded by the paucity of low-similarity models. But the 4% of models in the low-confidence distribution are distinctly worse than the other subsets. These observations suggest that AF2 produces models that are correct at the fold level more than 95% of the time, better than the previous two-thirds estimate [11].

### Discordance may limit model accuracy

The similar distribution between the high-confidence and the high-identity subsets (**Figure 1**) suggests that the main driver of the accuracy of most models is conformational change among multiple structures of nearly-identical sequences already in the PDB. Multiple PDB entries can reflect different conformations. Which protein conformation is more druggable depends on the clinical need associated with a particular disease state. AF2’s training algorithm handles multiple structure files in the PDB with the same primary sequence by down-selecting among them [11]. Structural details relevant for drug discovery, such as prosthetic groups, ion binding, and protons are not included, and AF2 models also exclude post-translational modifications.

An extreme conformational discordance case is calmodulin (**Figure 3**), a kinase that upon binding Ca^++^ ions changes from a globular to a dumb-bell shape by virtue of two hinge residues [24]. In the calmodulin AF2 model, the effects of discordance due to both conformations being present in the training set are reflected by a region of low-confidence scores at the hinge residues. Different AF2 runs yield slightly different results, but the resulting models split the difference between the two states (e.g., the AF2 model represented in grey on the top right side of **Figure 3**). This suggests that discordance among structures in the AF2 training set can result in composite models that may not necessarily accurately reflect the basic structure of any of the structures in the PDB, even for identical sequences.

**Figure 3.**
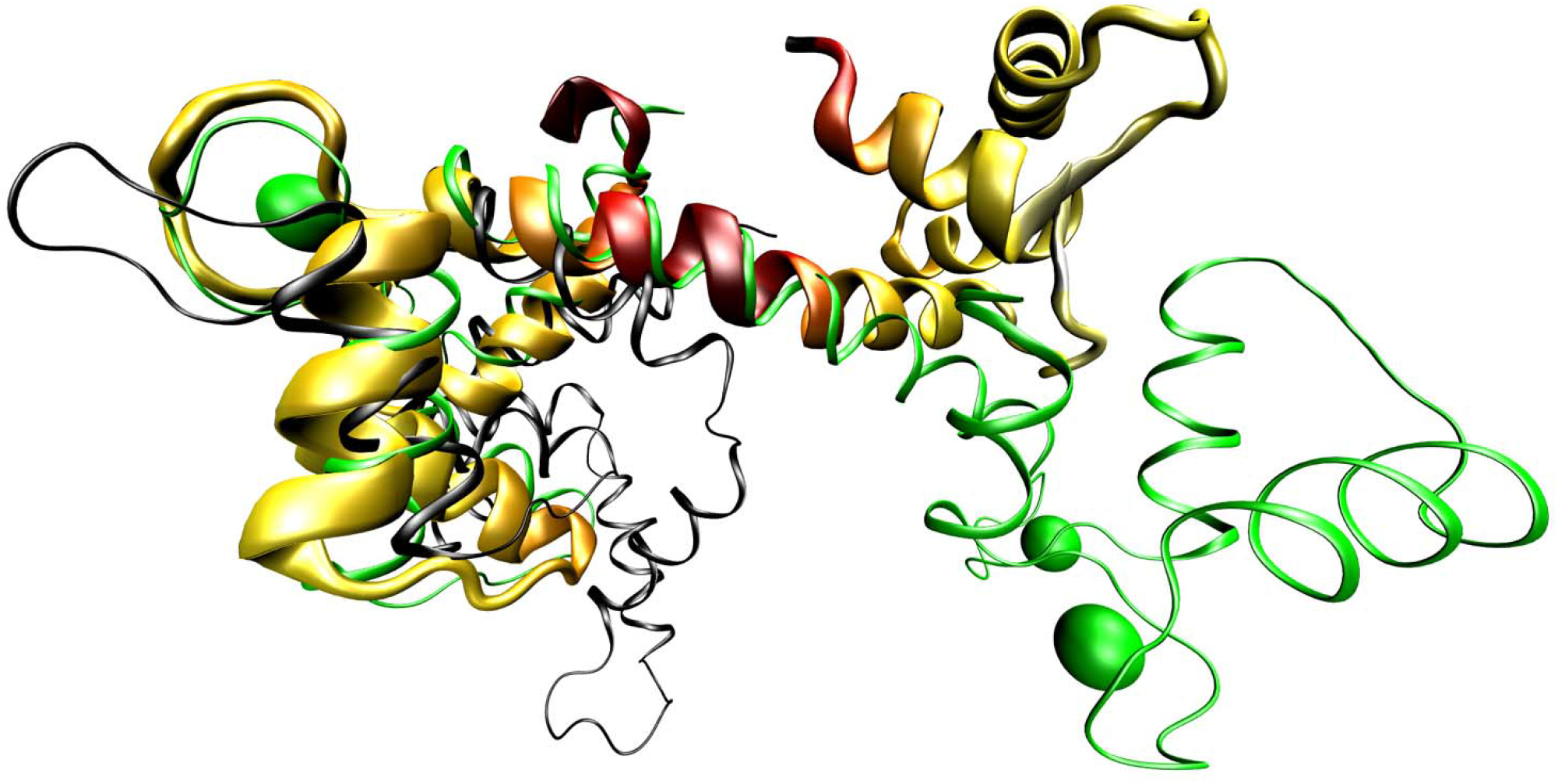
Comparison of discordant crystal structures of calmodulin with an AF model. The calcium-bound crystal structure (PDB entry 1CLL, thin green cartoon with Ca^++^ ions as spheres), with alignments against the first half of the calcium-free crystal structure (PDB entry 1QX5, thin black cartoon) and the AlphaFold2 model (P0DP23-F1-model_v1, thick yellow-red cartoon), aligned on their N-terminal halves. Yellow regions of the model represent very high confidence (pLDDT > 90) residues, while dark-red regions represent very low confidence (pLDDT<50) residues. The low confidence region at the center of the AF model corresponds to a hinge where the calcium-bound and calcium-free models diverge. When aligned in this manner, aRMSD values of 7.2 Å against the calcium-bound structure and 6.7 Å against the calcium-free structure were obtained. When aligned across all residues, the AF model yields aRMSDs of 10 Å against the calcium-bound structure and 17 Å against the calcium-free structure, respectively. Global LDDT scores for the experimental structures are 49% for all atoms and 56% for Cα only.

### Druggability: Are AF2 models ready for virtual screening?

Known protein conformational changes (as shown in **Figure 3**) can help us assess the effects of model accuracy on AF2 model readiness for target-based virtual screening (TBVS). If the model needs to be as close to the crystal structure as deoxy-myoglobin is to carboxy-myoglobin [25], only a tiny fraction of the AF2 models would be suitable for TBVS. If the model needs to be as close as R-state is to T-state hemoglobin, AF2 models may be suitable for characterizing allosteric sites [26]. A more typical TBVS example, where accuracy needs to be similar in capturing conformational changes, is when ERK2 is doubly phosphorylated. Given this example, a practical lower bound of global *pLDDT* of 80 could serve as basis for a model to likely be TBVS-ready. A value of *pLDDT* of 80 indicates a 68% likelihood of sidechain rotamers falling into the correct hemisphere (Fig 2B in [11]). Surfaces formed by two adjacent residues with *pLDDT* ≥ *80* are very close to the 50% accuracy limit if rotamer errors are independent. Having previously introduced *pLDDT*_*80*_, we set *pLDDT*_80_ ≥ 91.2 as criterion for assessing AF2 model quality, combined with the fraction of protein length for which this holds true (*pLDDT*_*80*__frac) to evaluate TBVS potential; see **Supplemental Information**. These criteria allow us to calculate a confusion matrix (see **Supplemental Figure S2)** that gives the sensitivity (true positive rate of classification) of 90.1% and a precision (positive predictive value of classification) of 86.3%.

Given these criteria, we evaluated which AF2 models of the human understudied proteins, Tdark [7], which currently lack an experimental PDB structure might be TBVS-ready (**Figure 4A)**. Of the set of 5592 “dark” proteins with AF2 models, 3051 (54.6%) meet our criteria for possibly being accurate enough for TBVS studies (**Figure 4B)**. Taking into account the estimated false-positive rate (∼6% of total), this implies that AF2 provides TBVS-ready models for about half of the understudied human proteins.

**Figure 4.**
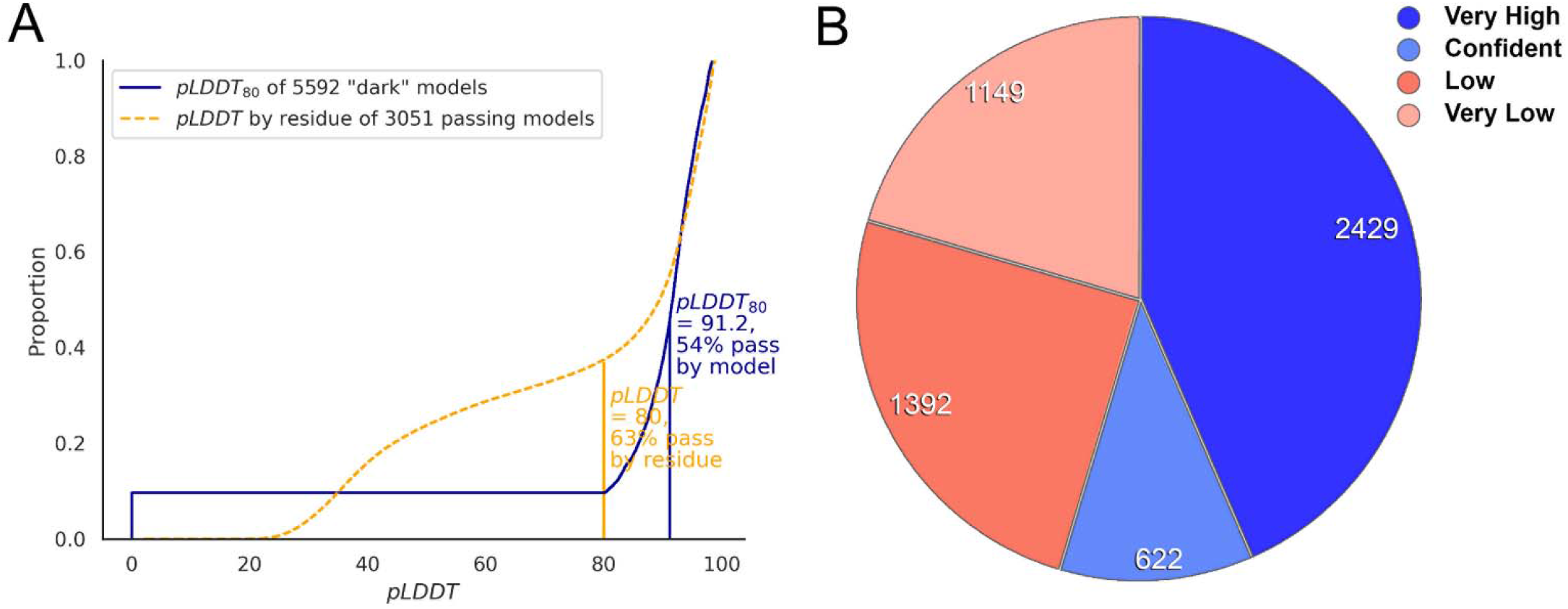
Fraction of the dark genome potentially illuminated by AF2 models. **A)** Of the set of 5592 unique “dark” proteins with AF2 models, 3051 (54%) pass the proposed selection criteria of pLDDT80 greater to or equal to 91.2 while having at least 20 residues with pLDDT >= 80. **B)** Pie chart illustrating AF2 model quality according to pLDDT80-derived criteria (see **Supplementary Information**): 3051 (54%) proteins associated with “very high” or “confident” AF2 models are likely to be TBVS-ready, whereas 2541 proteins are not.

## CONCLUSIONS

In our opinion, future work would do well to move away from the familiar aRMSD metric of overall model-structure agreement in favor of LDDT or other local measures. The aRMSD metric suggests that AF2 models are worse compared to pLDDT. Structural bioinformatics would also benefit from developing measures that disambiguate the effects of disorder, discord, and divergence.

AF2 forces us to reconsider the implications of disorder on druggability because it performs well at predicting IDRs [27]. Having trust that a region is disordered versus trusting the ordered region’s accuracy leads to different conclusions. Proteins containing IDRs play critical roles in many biological functions [28–35] and are associated with various diseases [36–39]. Thus, IDRs are potential targets in drug discovery [40–42]. “Disordered” does not mean “undruggable” because unique strategies for drug discovery in targets containing disordered regions are available [43]. Regions of pLDDT < 50 in an AF2 model indicate those strategies could be employed. Moreover, the existence of a structural model is neither necessary nor a sufficient condition for drug discovery. Even the use of high-quality experimental structures of the correct conformational state does not guarantee successful TBVS hits.

About 5% of the human “dark” proteome has structures in the PDB (Supplemental Information). Cost-to-benefit analyses of whether to deploy TBVS on AF2 models remain project-dependent. However, AF2 model quality may be “good enough” for rapid deployment for over 3000 understudied human proteins. AF2 models may help de-risk protein targets through protein expression and solubility and may provide protein engineering suggestions. By identifying likely boundaries of compact domains, disordered regions, or linkers, AF2 and other methods can enable synthesis-by-domain strategies that can break large proteins into more tractable modules to be expressed or synthesized then reconstituted in-vitro. Regardless of its impact on *in silico* technologies, AF2 does not preclude structural biology and structure-based drug design. However, AF2 is poised to become a powerful tool in the evolving drug discovery arsenal.

Computational models are very different from experimental structures in that they can be updated on-demand with the latest improvements. Public notebooks such as ColabFold [44] facilitate the removal of disordered termini, improving sequence alignment, adding a binding partner, and calculating new models within minutes. Although not designed with protein oligomers or assemblies in mind, multiple groups are working on use of AF2 to illuminate protein-protein interactions. In 2014, it was estimated that 40% of protein structures were experimentally determined [45]. With AF2 and future improvements, structural biology, and drug discovery are about to exponentially increase with new computational tools that combine sequence evolution, structures, and ligand binding knowledge [46].

## Supporting information

Excel file of classifications of dark protein

Tab-separated file of data on dark proteins, as produced by rafm

Tab-separated file of data on model/structure comparisons

Tab-separated file of per-residue data on files that pass the selection criterion

## Acknowledgements

We acknowledge support by the National Institutes for Health IDG KMC application from the University of New Mexico (Grant No. U24CA224370), the Passan Foundation (Grant No. R35 GM134864 to N.V.D.), the National Science Foundation (Grant No. 2137558 to Y.H.), and the National Science Foundation Graduate Research Fellowship (Grant No. DGE-1939267).

## Resources

We have implemented a *pLDDT*_80_ classifier along with other useful features as a python package called *rafm*, which is installable via the usual mechanism of “*pip install rafm”* at the command line on systems with python 3.8 or greater installed. https://github.com/unmtransinfo/rafm

**Supplementary Information** contains pLDDT and derivative score information for the PDB subset, and for the Tdark subset of the human proteome, as well as AF2 model evaluation criteria.

## Conflict of Interest

T.I.O. has received honoraria from or consulted for Abbott, AstraZeneca, Chiron, Genentech, Infinity Pharmaceuticals, Merz Pharmaceuticals, Merck Darmstadt, Mitsubishi Tanabe, Novartis, Ono Pharmaceuticals, Pfizer, Roche, Sanofi and Wyeth, and is on the Scientific Advisory Board of ChemDiv and InSilico Medicine.

---

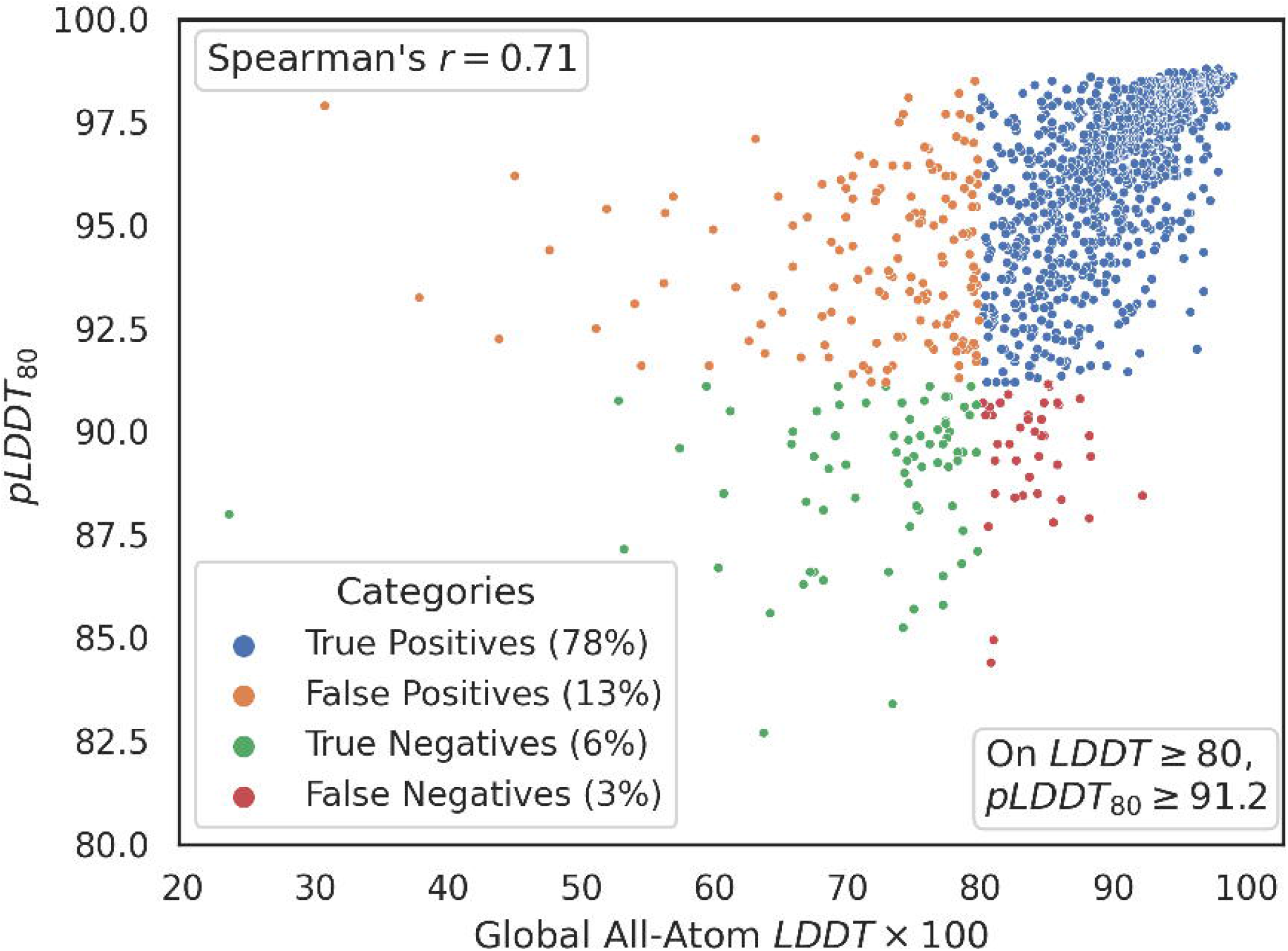

## Notes

https://bit.ly/reliableAF

