## Supplemental Information for "AlphaFold Models Illuminate Half of Dark Human Proteins"

**SUPPLEMENTAL MATERIAL:**

### The Post-AF2 Test Set

We started with a set of 2,604 crystal structures deposited in the PDB since AF was trained on April 30, 2018 until July 30, 2021, broken out by chain and limited to at least 10 residues in length. Pairwise alignments against UniProt sequences, using BLAST with default parameters, were made and consensus structural models using those alignments were constructed from AF and experimental data. In retrospect, these BLAST alignments were less than optimal for these purposes and future work should be done with a method for exact substring matching that would result in less rejection of likely-matching regions.

The comparison for intrinsically disordered regions (IDRs) was performed on N = 2604 proteins (Figure 1, main text).

For subsequent studies, the 2,604 proteins set was down-selected to not include any structures with residues that did not exactly match over the alignment (e.g., from expression tags that were included in the alignments). Out of the resulting N = 1810 proteins, an additional 31 protein models were excluded due to short sequences (up to 43 amino acids) representing less than 20% of the total protein, as captured by the AF2 models.

### Supplemental File Info

SupplementaryFile.xlsx

An Excel file with the two sheets, PDBvsAF2 and TdarkAF2. *For TdarkAF2, see below.*

**PDBvsAF2** has 1810 rows with the following columns:

1. PDB - Protein Data Bank Entry
2. Chain - chain from PDB entry
3. UniProt - UniProt ID of the protein corresponding to this chain
4. Length – number of residues in the protein associated with that UniProt ID
5. Model - name of the AF2 model file
6. Residues in LLDT - number of residues in the in the LDDT calculation, which is the number of residues in alignments less 1
7. LDDT - global all-atom LDDT
8. LDDT-Ca - global LDDT on C-alpha atoms only
9. RMSD - atomic RMSD on C-alpha atoms only, in Å
10. Residues in pLDDT - number of residues in the entire AF2 model
11. pLDDT_median – median value of pLDDT
12. pLDDT80_median - median value of pLDDT for those scores >= 80
13. pLDDT80_frac - fraction of the protein with pLDDT scores >= 80
14. Adj_pLDDT80frac - - fraction of pLDDT scores adjusted for the entire length of the protein
15. clustered-100% - true/false value indicating whether the model was a member of a cluster of all PDB sequences as downloaded on September 9, 2021 at 100% sequence identity over at least 80% of the length, using mmseqs version 13.4511 coverage mode 0 and cluster mode 2
16. clustered-5% - same as above, only with 5% sequence identity
17. AF2 Model quality - estimate of the model quality based on the following criteria:

| **Criteria** | **pLDDT80_median** | **Adj_pLDDT80fract** | **Count** |
| --- | --- | --- | --- |
| Very high | >=91.2 | >=0.5 | 839 |
| Confident | >=91.2 | 0.2 <= x < 0.5  *Or*  *x < 0.5 and residues_in_pLDDT >=100* | 655 |
| Low | < 91.2 | >= 0.2 | 166 |
| Very low | *any* | < 0.2 | 119 |
| Short sequence (*) | *Residues_in_LDDT < 45* | < 0.2 | 31 |

(*) These models were excluded from the discussion

All values computed on alignments between the experimental and those model residues with pLDDT > 50, except where noted.

**Validation of AF2 Model Quality Criteria:**

After excluding short sequence models, we compared “AF2 Model Quality” values (merging “very high” with “confident”, N = 1494; and “low” and “very low”, N = 285) with the number of protein models for which LDDT >= 0.8 (N = 1431) or LDDT < 0.8 (N = 348).

Our goal was to develop criteria for enrichment in models matching LDDT >= 0.8, that is “very high” and “confident” quality models, or “true positives” (TP). TP shows how many of the “good” quality models (LDDT >= 0.8) are correctly predicted by the “very high” or “confident” AF2 model quality criteria.

By the same token, true negatives (TN) shows how many models with LDDT < 0.8 values are correctly labeled “(very) low” by the “(very) low” AF2 model quality labels.

**Confusion Matrix	Count**

True positives (TP)	1289

False positives (FP)	205

False negatives (FN)	142

True negatives (TN)	143

From the above values, we then calculated

(1) Sensitivity, or true positive rate = 0.9008

TPR = TP (TP + FN)

(2) Precision, or positive predictive value = 0.8628

PPV = TP (TP + FP)

(3) Specificity, SPC = 0.4109

SPC = TN (TN + FP)

The above criteria indicate that the AF2 model quality criteria correctly identify 90% of the “good” (LDDT >= 0.8) models, and that 86.3% of the “very high” and “confident” models are likely to be of “good” (cf LDDT) quality. However, the criteria lack specificity, i.e., they do not eliminate “poor” (cf LDDT) models. This behavior is in alignment with our conclusion that “pLDDT confidence scores below 80 are more indicative of disorder than of confidence in the accuracy of ordered structures”

**TdarkAF2** contains a list of 5991 understudied human proteins which are classified as “Tdark” according to Target Development Levels described in Oprea et al., Nature Rev Drug Discov 2018.

Of these, N = 299 proteins are associated with at least 1 PDB file (yellow background indicates proteins associated with ligands in PDB as well). Data is current as of July 2021. For another N = 100 proteins, AF2 failed to generate a model.

For the remaining 5592 proteins we provide per-model statistics of AF2 models as well as AF2 Model Quality criteria assessment. The columns are as follows:

1. UniProt – UniProt ID of the protein
2. Length – number of residues in the protein associated with that UniProt ID
3. Model – the name of the specific AF2 model evaluated (some proteins have multiple models; the model with the highest number of residues and pLDDT80_median is shown)
4. residues_in_pLDDT – number of residues in the model
5. pLDDT_median – median value of pLDDT
6. pLDDT80_median – median value of those pLDDT scores above 80 over the entire model
7. pLDDT80_frac – fraction of residues in the model with pLDDT scores above 80
8. Adj_pLDDT80frac - - fraction of pLDDT scores adjusted for the entire length of the protein
9. AF2 Model quality - estimate of the model quality based on the following criteria:

| **Criteria** | **pLDDT80_median** | **Adj_pLDDT80fract** | **Count** |
| --- | --- | --- | --- |
| Very high | >=91.2 | >=0.5 | 2429 |
| Confident | >=91.2 | 0.2 <= x < 0.5  *Or*  *x < 0.5 and residues_in_pLDDT >=100* | 622 |
| Low | < 91.2 | >= 0.2 | 1392 |
| Very low | *any* | < 0.2 | 1149 |

### Supplemental Figures

**Supplemental Figure S1.** **Scatterplot of median pLDDT (top) and pLDDT_80_ (bottom) against log(Cα-aRMSD).** Changing the measure from median *pLDDT* to median of only those scores greater than 80 (*pLDDT_80_*) improves the Spearman’s Rank correlation coefficient on *aRMSD* from -0.37 (top) to -0.48 (bottom). Blue dots represent models in low-identity (5% sequence identity) clusters.

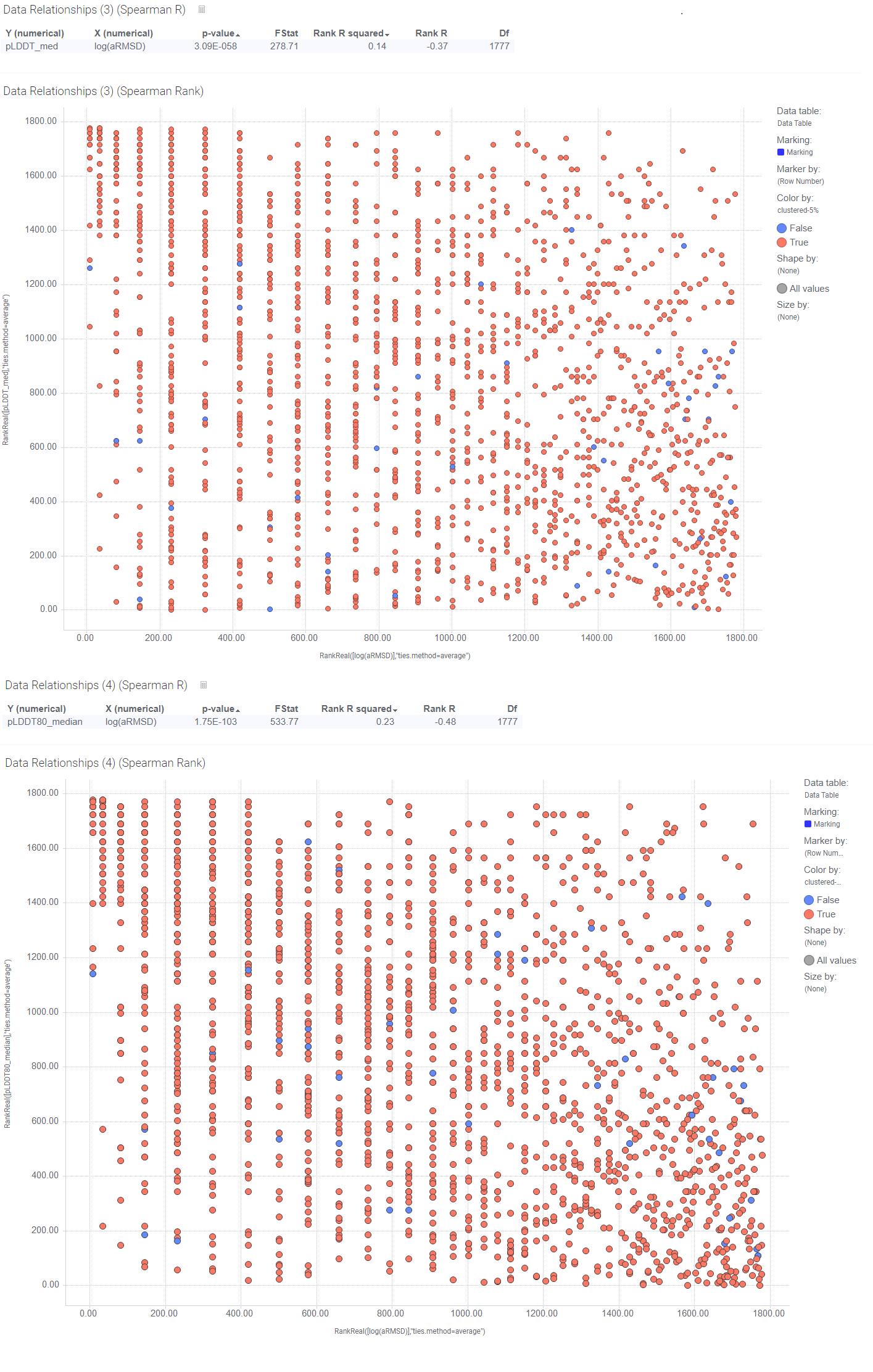

**Supplemental Figure S2. Scatter plot of pLDDT_80_ against global all-atom LDDT.** Using an value of *LDDT* greater than or equal to 80 as the measure of usability on our set of 1779 downselected post-AF2 crystal structures gives 1289 models as true positives, 205 models as false positives, 142 models as false negatives, and 143 models as true negatives. Our classifier therefore has a sensitivity (true positive rate) of 90.08% and a precision (positive predictive value) of 86.28%. Both are indicative of a good performance in enrichment for high-quality AF2 models.

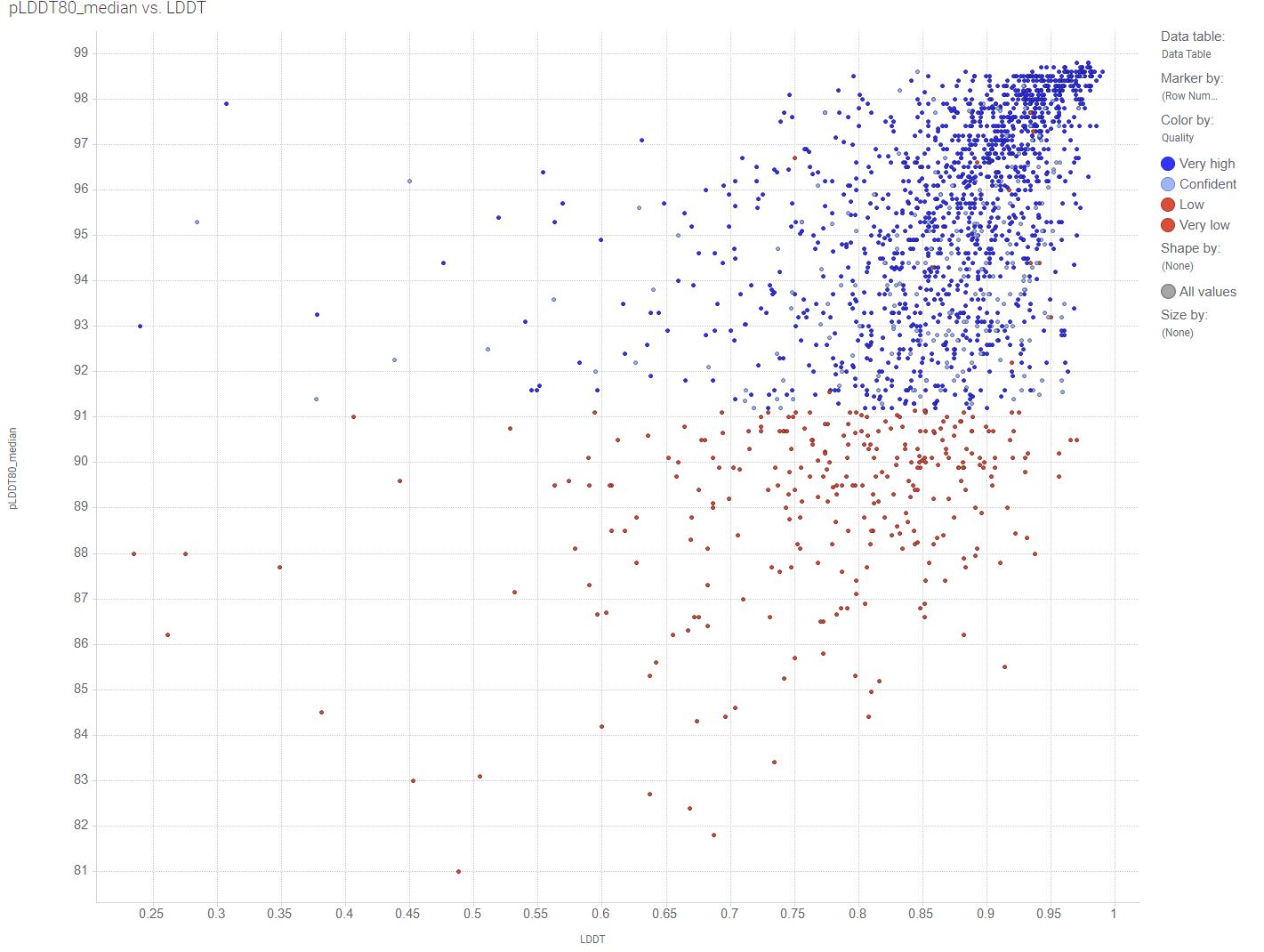
